# Intra-mitochondrial proteostasis is directly coupled to alpha-synuclein and Amyloid β 1-42 pathology

**DOI:** 10.1101/561134

**Authors:** Janin Lautenschläger, Sara Wagner-Valladolid, Amberley D. Stephens, Ana Fernández-Villegas, Colin Hockings, Ajay Mishra, James D. Manton, Marcus J. Fantham, Meng Lu, Eric J. Rees, Clemens F. Kaminski, Gabriele S. Kaminski Schierle

## Abstract

Mitochondria have long been implicated in Parkinson’s disease (PD), however, it is not clear how mitochondrial impairment and alpha-synuclein pathology are coupled. We report here that intra-mitochondrial protein homeostasis plays a major role in alpha-synuclein fibril elongation, as interference with intra-mitochondrial proteases and mitochondrial protein import significantly aggravate alpha-synuclein aggregation. In contrast, direct inhibition of mitochondrial complex I, increase in intracellular calcium concentration or formation of reactive oxygen species (ROS), all of which have been associated with mitochondrial stress, did not affect alpha-synuclein pathology. We further demonstrate that similar mechanisms are involved in Amyloid β 1-42 (Aβ42) aggregation, suggesting that mitochondria are directly capable of influencing cytosolic protein homeostasis of aggregation-prone proteins.

## Introduction

Parkinson’s disease (PD) is the second most common neurodegenerative disease and affects about 1% of the population over 60 years (1). Alpha-synuclein aggregation has been found central to the disease, since SNCA mutations are associated with familiar PD (2) and alpha-synuclein has been identified as a major constituent of Lewy bodies in sporadic PD and dementia with Lewy bodies (3). A relationship between protein aggregation and protein levels is demonstrated by familiar PD with SNCA gene duplication and triplication (4–6), however the trigger for protein aggregation in sporadic cases is less clear. Likewise, also Alzheimer’s disease is linked to increased protein aggregation, where enhanced levels of Amyloid beta (Aβ) due to mutations in the genes coding for the amyloid precursor protein (APP) (7, 8) or presenilin (9–11) are found in familiar AD.

Mitochondria have long been implicated in PD, ever since the discovery that inhibitors of the mitochondrial complex I can lead to dopaminergic neuron death (12–16). But also the regulation of mitophagy via the PTEN-induced kinase 1 (PINK1) plays a role in PD (17) and seems to be coupled to alpha-synuclein toxicity. PINK1 overexpression is able to decrease the effect of alpha-synuclein toxicity in Drosophila (18, 19), and PINK1 knockout in mice increases alpha-synuclein neurotoxicity (20, 21). Furthermore, PINK1 patient iPSC-derived midbrain dopaminergic neurons show accumulation and aggregation of alpha-synuclein (Chung *et al*, 2016), and PINK1 knockout rats even show alpha-synuclein de novo aggregation (24).

We have demonstrated previously that alpha-synuclein interaction with calcium leads to conformational changes at the C-terminus of alpha-synuclein, but also at the aggregation-prone non-amyloid component (NAC)-region, suggesting that calcium can directly influence the aggregation propensity of alpha-synuclein (25). Thus, we tested, whether treatment with BAPTA-AM, which is supposed to decrease intracellular calcium by calcium chelation, was able to decrease alpha-synuclein pathology. However, we observed drastically enhanced alpha-synuclein aggregation, which we could attribute to mitochondrial fragmentation. This led us to study and show that disturbances in intra-mitochondrial proteostasis can aggravate alpha-synuclein aggregation. We identified that the Lon protease and the high-temperature requirement protein A2 (HtrA2) protease, as well as mitochondrial protein import are crucial in determining the level of alpha-synuclein seeding in a cellular model. However, inhibition of the mitochondrial complex I and a direct increase in cytosolic calcium or oxidative stress were not able to reproduce enhanced alpha-synuclein pathology such as observed in cells upon inhibition of mitochondrial protein homeostasis. In addition, the mitochondrial protease HtrA2 and mitochondrial protein import were also relevant for Aβ42 aggregation and isolated mitochondria were shown to directly diminish Aβ42 aggregation.

## Results

### BAPTA-AM increases alpha-synuclein pathology

SH-SY5Y cells overexpressing YFP-alpha-synuclein were incubated for 4 hours (h) with small fibrillar seeds made of unlabelled human recombinant alpha-synuclein to study alpha-synuclein pathology after seeding, as described previously (26–30). Cells were left in culture for 3 days before the level of alpha-synuclein aggregation within the cells was determined (see Supplementary Fig. 1A and B for treatment regime and fibrillar seeds). While unseeded YFP-alpha-synuclein overexpressing cells did not display any aggregates, those that were seeded displayed large YFP-alpha-synuclein aggregates which were built up from fine filaments (Supplementary Fig. 1C). Furthermore, YFP-alpha-synuclein inclusions stained positive for ubiquitin and p62, which are both characteristic markers of Lewy bodies in human disease (26) (Supplementary Fig. 1D and E).

We have previously shown that alpha-synuclein strongly interacts with calcium, leading to conformational changes both at the C-terminal calcium-binding domain, and the aggregation-prone non-amyloid component (NAC)-region, which suggests that calcium can directly influence the aggregation propensity of alpha-synuclein. Consistently, increased calcium concentrations significantly enhanced alpha-synuclein nucleation in vitro (25). BAPTA-AM as a calcium chelator is supposed to decrease cytosolic calcium and has been previously reported to alleviate KCl-induced alpha-synuclein aggregation (31). However, when we treated the above-described cells with BAPTA-AM before the incubation with fibrillary seeds (1h) or before and during incubation with fibrillary seeds (5h) alpha-synuclein seeding was drastically increased (Fig. 1A). We thus tested the calcium effect of BAPTA-AM in SH-SY5Y cells and verified that BAPTA-AM was able to decrease cytosolic calcium. However, the effect was only transient and calcium chelation was already compensated in cells treated for 1 h and 5 h (Fig. 1B, fluorescence lifetime decrease after 10 min from 2381+/-8 ps to 2170+/-15 ps, lifetime of 2400 +/-8 ps after 1h and 2460+/-12 ps after 5h). Since the 1 h treatment showed calcium levels comparable to control, but already increased alpha-synuclein aggregation, suggested that the increase of alpha-synuclein seeding by BAPTA-AM was not mediated via calcium. In addition, we tested whether both the ester form of BAPTA, BAPTA-AM, as well as the active BAPTA itself affected the aggregation of alpha-synuclein directly, but found no difference in alpha-synuclein aggregation kinetics measured in-vitro by Thioflavin T (ThT) fluorescence (Fig. 1C, t50 125.6+/-8.6 h and 122.6+/-7.2 h vs. 116.6+/− 11.1 h). This confirms that the effect of BAPTA is most likely triggered by a cellular response. In the literature there was one report describing that BAPTA-AM can lead to mitochondrial fragmentation (32), so we stained our cells with Mitochondria-RFP and indeed observed a drastic mitochondrial fragmentation upon BAPTA-AM treatment (Fig. 1D).

**Fig. 1.**
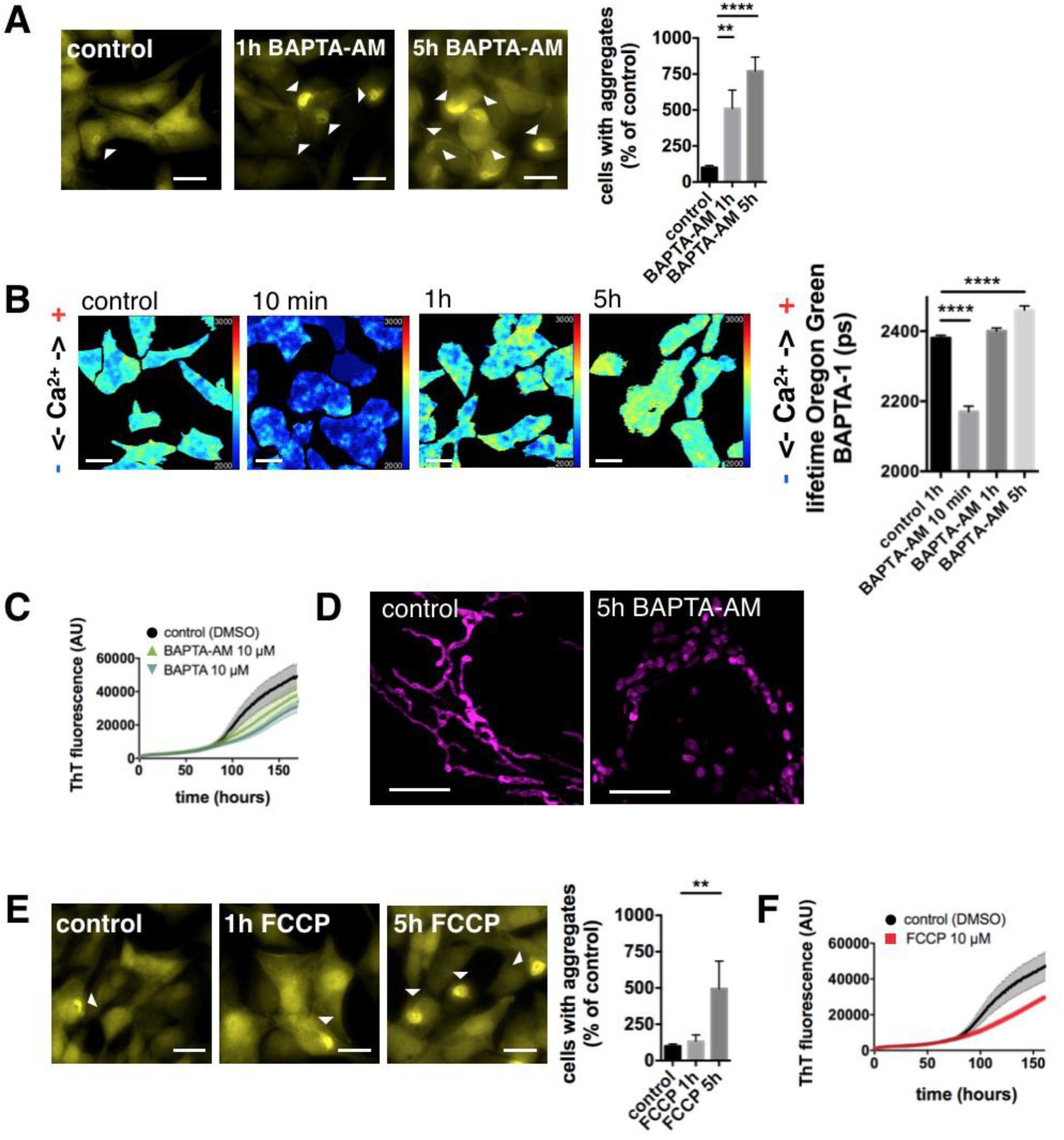
BAPTA-AM increases alpha-synuclein pathology. (A) YFP-alpha-synuclein SH-SY5Y cells were treated with DMSO (control), 10 µM BAPTA-AM for 1h (before fibrillar seed incubation) and for 5h (before plus during the incubation with alpha-synuclein fibrillar seeds). Scale bars: 20 µm. Alpha-synuclein seeding was increased upon 1h pre-treatment and 5h treatment with BAPTA-AM. Data are presented as mean ± SEM. *p = 0.0127 and ****p < 0.0001 (Kruskal-Wallis test with Dunn’s multiple comparison). N = 16, 9, 15 with n = regions analyzed, three biological repeats. (B) Fluorescence lifetime images of cytosolic calcium levels (Oregon Green^TM^ 488 BAPTA-1 fluorescence lifetime) in SH-SY5Y cells treated with DMSO (control), 10 µM BAPTA-AM for 10 min, 1h or 5h. The cytosolic calcium level within cells was significantly reduced upon 10 min incubation with BAPTA-AM, however after 1h of incubation with BAPTA-AM calcium levels returned back to basal levels. After 5h treatment with BAPTA-AM calcium levels significantly increased beyond basal calcium levels. Data are presented as mean ± SEM. ****p < 0.0001 (Kruskal-Wallis test with Dunn’s multiple comparison). N = 88, 54, 61, 46, with n = cells analyzed, three biological repeats. (C) ThT assay displaying the aggregation kinetics of alpha-synuclein in-vitro in the presence of DMSO, 10 µM BAPTA-AM, or 10 µM BAPTA. Data are presented from three biological repeats. (D) Mito-RFP stained mitochondrial network in SH-SH5Y cells. Cells were treated with DMSO (control) or 10 µM BAPTA-AM for 5h. Scale bars: 5 µm. (E) YFP-alpha-synuclein SH-SY5Y cells treated with DMSO (control), 10 µM FCCP for 1h (before fibrillar seed incubation) and for 5h (before plus during the incubation with alpha-synuclein fibrillar seeds). Scale bars: 20 µm. Alpha-synuclein seeding was increased upon 5h treatment with FCCP. Data are presented as mean ± SEM. **p = 0.0064 (Kruskal-Wallis test with Dunn’s multiple comparison). N = 16, 6, 9 with n = regions analyzed, three biological repeats. (F) ThT assay displaying the aggregation kinetics of alpha-synuclein in-vitro in the presence of DMSO or 10 µM FCCP. Data are presented from three biological repeats.

Thus, it seems that mitochondrial dysfunction influences alpha-synuclein aggregation per se, which we tested using carbonyl cyanide 4-(trifluoromethoxy)phenylhydrazone (FCCP), a mitochondrial uncoupler which dissipates the mitochondrial membrane potential. Treatment with FCCP during alpha-synuclein fibril incubation (5h) increased alpha-synuclein seeding (Fig. 1E), however FCCP did not increase alpha-synuclein aggregation in-vitro (t50 117.0+/-9.8 h vs. 115.6+/-10.1 h), confirming that its effect in cells is the result of a cellular response rather than a direct interaction of FCCP and alpha-synuclein (Fig.1F).

### Classical downstream effectors of mitochondrial dysfunction are unable to influence alpha-synuclein pathology

We next tested whether downstream events of mitochondrial dysfunction could reproduce increased alpha-synuclein seeding. However, if we inhibited mitochondrial ATP production via inhibition of complex I, directly increased cytosolic calcium or oxidative stress for 3 days until seeding was evaluated, no increase in alpha-synuclein seeding was seen (Fig. 2A). We used 1-methyl-4-phenylpyridinium (MPP^+^), the active metabolite of 1-methyl-4-phenyl-1,2,3,6-tetrahydropyridine (MPTP) to inhibit complex I of the electron transport chain, ionomycin, an ionophore, to increase cytosolic calcium concentrations via calcium influx through the plasma membrane, and menadione to induce the formation of ROS via redox-cycling (33). To test that the inhibitors were active we measured ATP, calcium and H2O2 in SH-SY5Y cells using the fluorescent sensors Ateam1.03, Oregon-Green^TM^ BAPTA-1 and HyPer. The readout of the fluorescence lifetime of these sensors allows to estimate and directly compare the different treatments (34, 35). So, MPP^+^ inhibition of complex I reduced the ATP level (Fig. 2B, fluorescence lifetime increase of the FRET donor from 1298 +/− 17 ps to 1511 +/− 20 ps), ionomycin increased cytosolic calcium (Fig. 2C, fluorescence lifetime increase of Oregon-Green^TM^ BAPTA-1 from 2381 +/− 8 ps to 2663 +/− 7 ps), and menadione increased H2O2 levels (Fig. 2D, fluorescence lifetime decrease of cpYFP from 1575 +/− 4 ps to 1557 +/− 3 ps). Treatment with MPP^+^ lead to lower ATP depletion than under FCCP, but to higher ATP depletion than under BAPTA-AM (Fig. 2B). Ionomycin treatment lead to higher calcium increase compared to both, FCCP and BAPTA-AM (Fig 2C). And menadione treatment lead to comparable levels of H2O2 increase than under treatment with FCCP and BAPTA-AM (Fig. 2D). This suggests that complex I inhibition, cytosolic calcium increase and oxidative stress do not aggravate alpha-synuclein pathology per se.

**Fig. 2.**
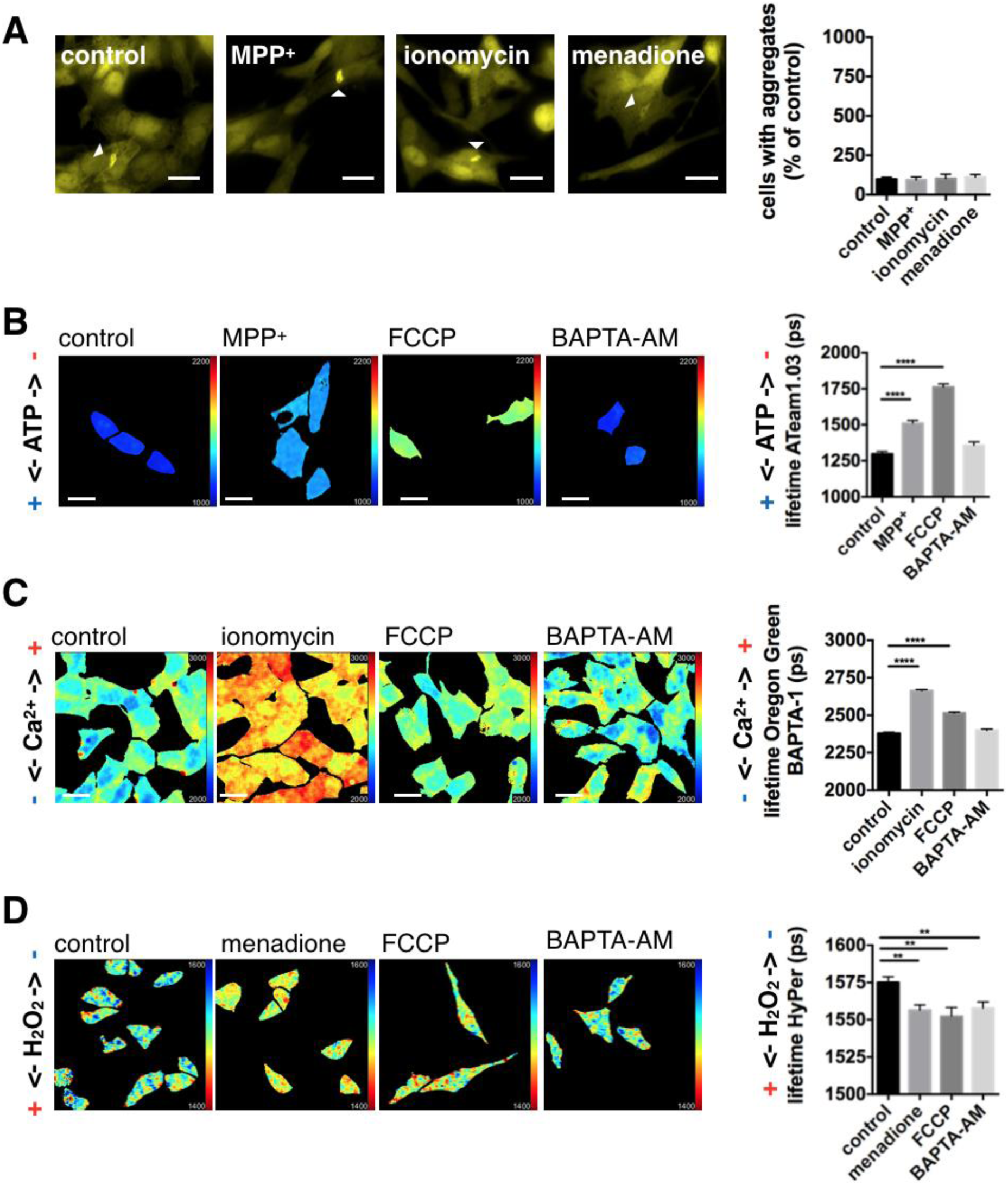
Downstream effectors of mitochondrial dysfunction do not influence alpha-synuclein pathology. (A) YFP-alpha-synuclein SH-SY5Y cells were treated with DMSO (control), 500 µM MPP^+^, 1 µM ionomycin, or 3 µM menadione for 3 days (1h before, during alpha-synuclein fibrillar seed incubation, and during the 3 day period until evaluation). Scale bars: 20 µm. Alpha-synuclein seeding was not significantly increased (one-way ANOVA with Dunnett’s post-hoc correction). Data are presented as mean ± SEM, N = 11, 8, 8, 7 with n = regions analyzed, three biological repeats. (B) Fluorescence lifetime images and graphs for ATP levels (Ateam1.03 fluorescence lifetime) in SH-SY5Y cells treated with DMSO (control), 500 µM MPP^+^, 10 µM FCCP and 10 µM BAPTA-AM for 1h. The effect of 500 µM MPP^+^ and FCCP was significant, BAPTA-AM had no significant effect. ****p < 0.0001 and N = 43, 74, 48, 47. (C) Fluorescence lifetime images and graphs for cytosolic calcium levels (Oregon Green^TM^ 488 BAPTA-1 fluorescence lifetime) in SH-SY5Y cells treated with DMSO (control), 1 µM ionomycin, 10 µM FCCP and 10 µM BAPTA-AM for 1h. The effect of 1 µM ionomycin and FCCP was significant. Both, the effect of FCCP and BAPTA-AM treatment were lower than seen with ionomycin. ****p < 0.0001 and N = 88, 60, 42, 61. (D) Fluorescence lifetime images and graphs of H2O2 levels (HyPer fluorescence lifetime) in SH-SY5Y cells treated with DMSO (control), 3 µM menadione, 10 µM FCCP and 10 µM BAPTA-AM for 1h. The effect of 3 µM menadione, FCCP and BAPTA-AM was significant. Both, FCCP and BAPTA-AM did not produce a higher effect than seen with menadione. **p = 0.0017, p = 0.0012 and p = 0.0058 and N = 79, 63, 36, 70. All scale bars: 20 µm. Data are presented as mean ± SEM with n = cells analyzed, Kruskal-Wallis test with Dunn’s multiple comparison, three biological repeats.

### Inhibition of mitochondrial proteostasis increases alpha-synuclein pathology

In the next step we evaluated mitochondrial fragmentation upon treatment with FCCP and BAPTA-AM, as well as MPP^+^, ionomycin and menadione. Automated analysis of mitochondrial length showed that both FCCP and BAPTA-AM treatment leads to mitochondrial fragmentation, while neither MPP^+^, nor ionomycin or menadione influenced mitochondrial integrity (Fig. 3A and B). Furthermore, we saw that FCCP treatment led to higher levels of mitochondrial fragmentation compared to BAPTA-AM, though the effect of BAPTA-AM on alpha-synuclein seeding was more pronounced than after FCCP treatment (refer to Fig. 1A and E). This indicated that mitochondrial fragmentation is not directly linked to alpha-synuclein aggregation and additional factors might play a role.

**Fig. 3.**
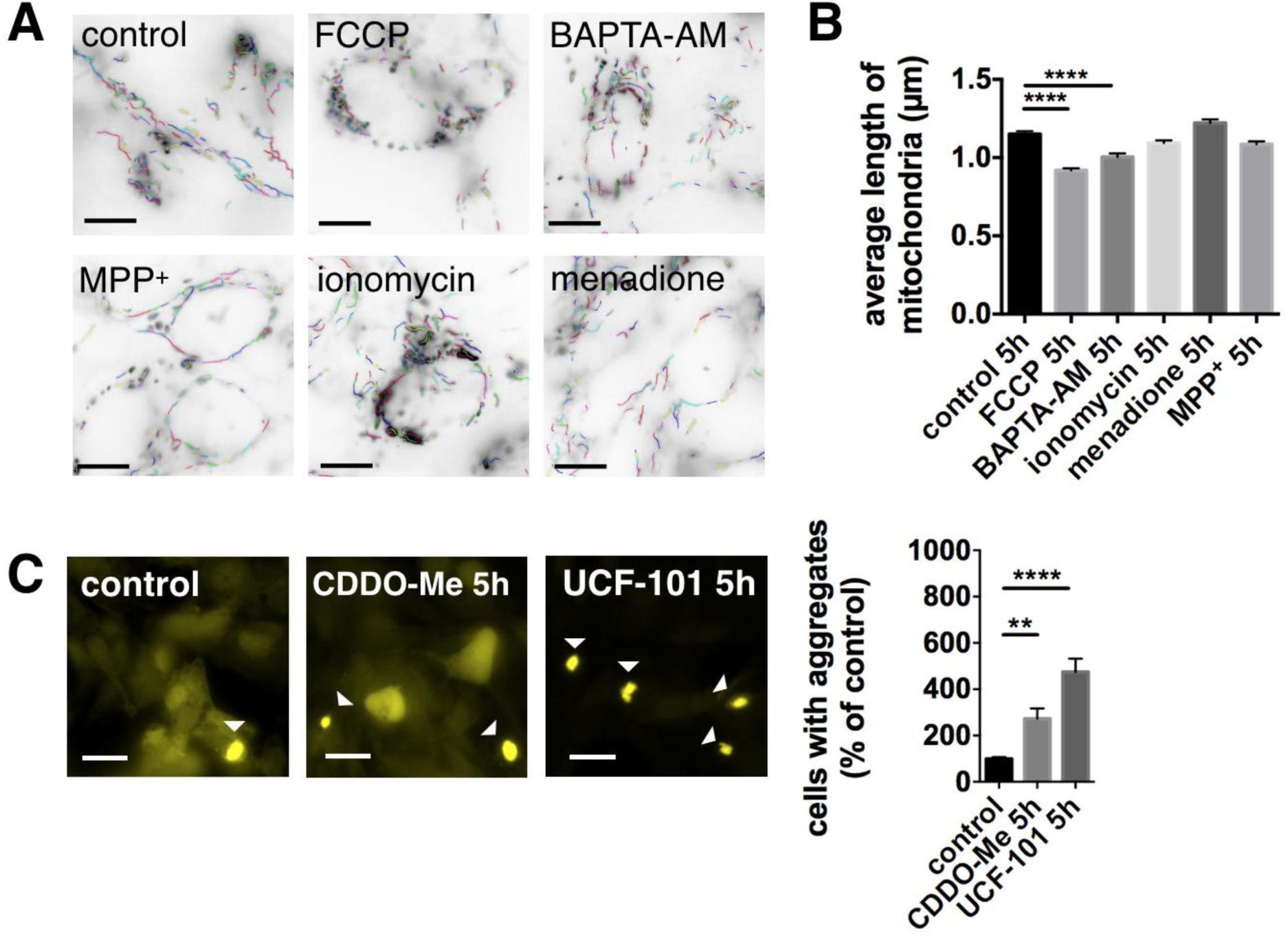
Inhibition of mitochondrial proteases increases alpha-synuclein pathology. (A and B) Quantification of mitochondrial fragmentation after 5h treatment with 10 µM FCCP, 10 µM BAPTA-AM, 500 µM MPP+, 1 µM ionomycin and 3 µM menadione. The mitochondrial length was significantly decreased after treatment with FCCP and BAPTA-AM. Scale bars: 10 µm. Data are presented as mean ± SEM. ****p < 0.0001 (Kruskal-Wallis test with Dunn’s multiple comparison). N = 76, 92, 103, 90, 88, 89 with n = individual images, three biological repeats. Images analysis of mitochondrial fragmentation was performed using NIEL Mito (86), (C) YFP-alpha-synuclein SH-SY5Y cells were treated with DMSO (control), 1 µM CDDO-Me, or 20 µM UCF-101 before and during the incubation with alpha-synuclein fibrillar seeds (5h). Scale bars: 20 µm. Alpha-synuclein seeding was increased upon both treatments. Data are presented as mean ± SEM. **p = 0.005 and ****p < 0.0001 (Kruskal-Wallis test with Dunn’s multiple comparison). N = 15, 9, 11 with n = regions analyzed, three biological repeats.

Previously, BAPTA-AM has been reported to inhibit proteases (36–38), which is mediated via the blocking of intracellular calcium transients required to regulate protease activity (39, 40). This led us to test the effect of mitochondrial proteostasis on alpha-synuclein pathology. We treated the YFP-alpha-synuclein SH-SY5Y cells with CDDO-Me, an inhibitor of the Lon protease (41), which has recently been shown to influenced aggregate dissolution after heat shock (42) and also with UCF-101, an inhibitor of the high temperature requirement protein A2 (HtrA2/Omi) protease (43), which has previously been linked to PD. Our results show that both mitochondrial protease inhibitors significantly increased alpha-synuclein pathology (Fig. 3C), where the effect of HtrA2 protease inhibition was superior to Lon protease inhibition.

### Inhibition of mitochondrial proteostasis increases Amyloid β 1-42 pathology

To test if the above-discussed mechanisms also contribute to the aggregation of other proteins involved in neurodegeneration, we investigated the effect of mitochondrial proteostasis on Amyloid β 1-42 (Aβ42) pathology. We used a stable HEK cell line overexpressing Aβ42-mCherry via a tetracycline-inducible expression system as published previously (44). After induction of Aβ42-mCherry expression, the cells were treated with FCCP, BAPTA-AM and the protease inhibitors as in the alpha-synuclein aggregation model. We found that both, FCCP and BAPTA-AM, increased the aggregation of Aβ42 (Fig. 4A). BAPTA-AM had a more pronounced effect compared to FCCP, similar to what has been seen for alpha-synuclein aggregation. Inhibition of the Lon protease did not significantly increase Aβ42 aggregation, however inhibition of HtrA2 using UCF-101 increased Aβ42 pathology (Fig. 4B). Vice versa, enhancing mitochondrial proteostasis via HtrA2 overexpression decreased Aβ42 aggregation, demonstrating that mitochondrial proteostasis plays a role in regulating the aggregation of amyloidogenic proteins (Fig. 4C).

**Fig. 4.**
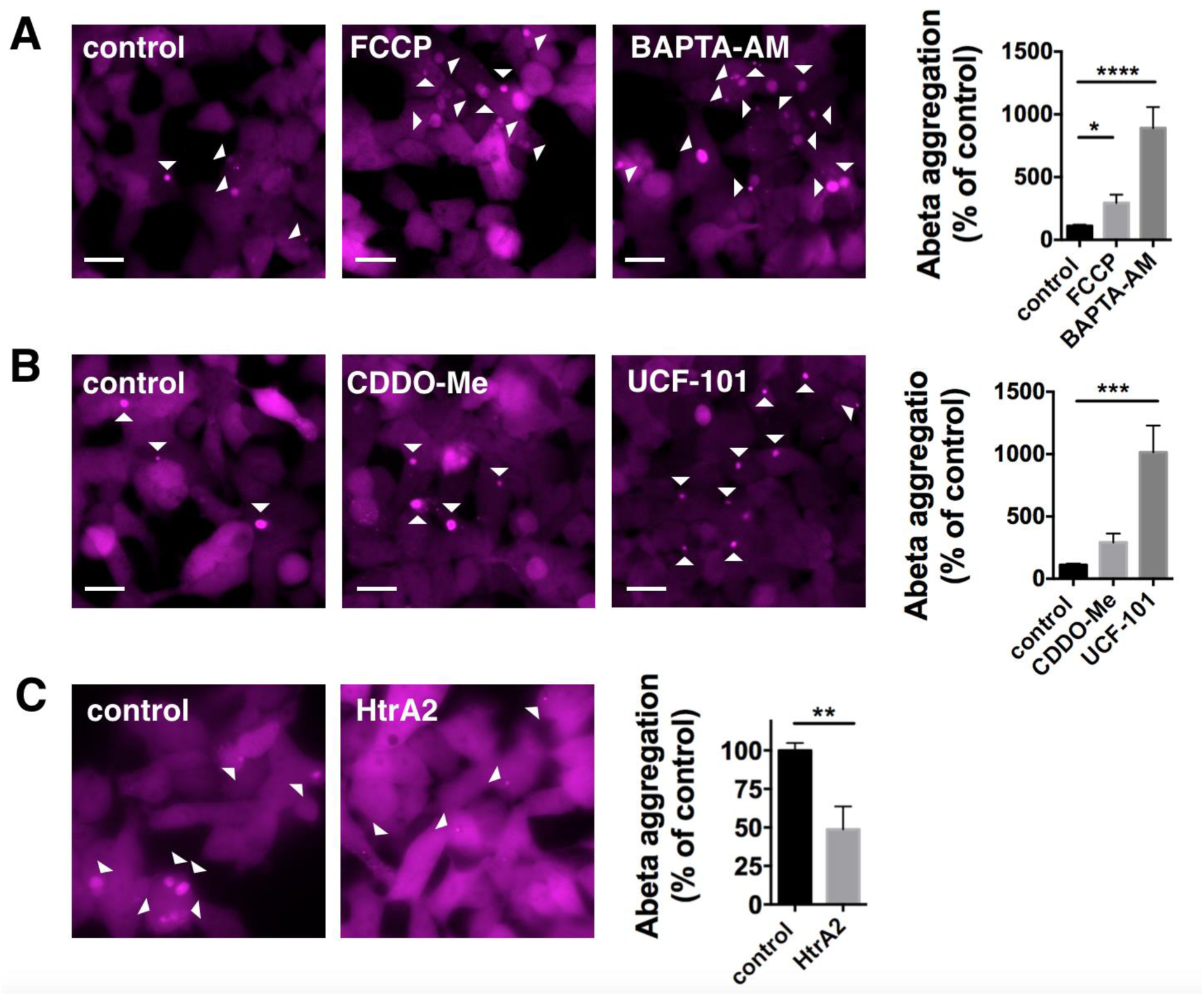
Mitochondrial proteostasis influences Amyloid β 1-42 pathology. (A) Aβ42-mCherry overexpressing HEK cells were treated with DMSO (control), 1 µM FCCP or 10 µM BAPTA-AM for 24 h. The aggregation of Aβ42 was increased upon treatment with FCCP and BAPTA-AM. Data are presented as mean ± SEM. *p = 0.0298 and ****p < 0.0001 (Kruskal-Wallis test with Dunn’s multiple comparison). N = 9, for all conditions, with n = wells analyzed, three biological repeats. (B) Aβ42-mCherry cells were treated with DMSO (control), 0.1 µM CDDO-Me or 20 µM UCF-101 for 24 h. The aggregation of Aβ42 was increased upon treatment with UCF-101. Data are presented as mean ± SEM. ***p = 0.0001 (one-way ANOVA with Dunnett’s post-hoc correction). N = 9, for all conditions, with n = wells analyzed, three biological repeats. (C) Aβ42-mCherry cells were transfected with either uncut pcDNA3 (control) or HtrA2 pcDNA3 and Aβ42-mCherry expression was induced with tetracycline for 3 days. The aggregation of Aβ42 was significantly decreased upon overexpression of HtrA2. Data are presented as mean ± SEM. **p = 0.0089 (two-tailed unpaired t-test). N = 9, 9 with n = regions analyzed, three biological repeats. Scale bars: 20 µm.

### In-vitro aggregation of Amyloid β 1-42 is influenced by mitochondria and HtrA2

In order to show that mitochondria directly influence Aβ42 proteostasis, we analyzed Aβ42 aggregation in vitro using a fluorescence lifetime aggregation assay. This assay makes use of the reduction in fluorescence lifetime of labelled proteins when they start to aggregate and are tightly packed, as described in detail by our group before (45, 46). We used Aβ42 containing 50 % Hylite^TM^ Fluor 488 labelled Aβ42 which was incubated for 2h at room temperature, showing a reduction of Hylite^TM^ Fluor 488 fluorescence lifetime from 3380 +/− 93 ps to 3003 +/− 97 ps within this time interval (Fig. 5A control t0 and t2h). However, in the presence of isolated rat brain mitochondria, only a non-significant drop in Aβ42 Hylite^TM^ Fluor 488 fluorescence lifetime occurred (Fig. 5A mito t0 and t2h, 3538 +/− 15 ps compared to 3502 +/− 5 ps). Note, the fluorescence lifetime of Aβ42 Hylite^TM^ Fluor 488 incubated with mitochondria is higher at the beginning of the experiment (t0) than in the control group (Aβ42 + mito 3538 +/− 15 ps compared to Aβ42 control with 3380 +/− 93 ps), since in the control Aβ42 starts to aggregate immediately upon preparation. The decrease in Aβ42 Hylite^TM^ Fluor 488 fluorescence lifetime seen in the presence of mitochondria was only minor, however, if we pre-incubated mitochondria with UCF-101 Aβ42 Hylite^TM^ Fluor 488 fluorescence lifetime significantly decreased over the 2 h time interval, demonstrating that Aβ42 aggregation is increased upon inhibition of HtrA2 (Fig. 5B, UCF-101 t0 and t2h, 3523 +/− 16 ps compared to 3429 +/− 20 ps).

**Fig. 5.**
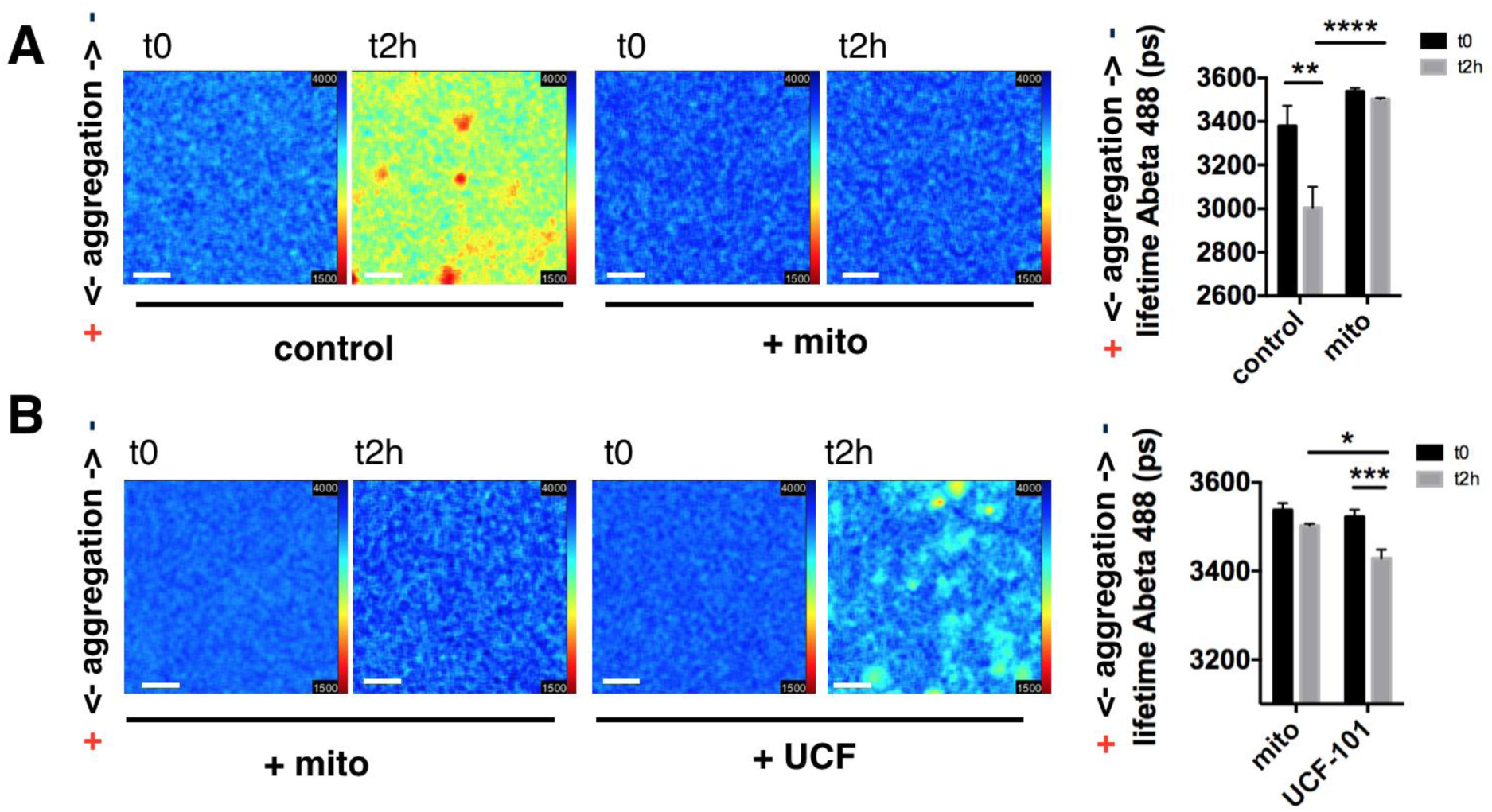
HtrA2 influences in-vitro aggregation of Aβ42. (A) Fluorescence lifetime images of Hilyte™ Fluor 488 labelled Aβ42 at the beginning of the experiment (t0) and after 2 h of incubation at room temperature (t2h) show a decrease in fluorescence lifetime in control conditions demonstrating protein aggregation. No decrease in Aβ42 Hilyte™ Fluor 488 fluorescence lifetime was seen upon addition of isolated mitochondria. Data are presented as mean ± SEM. **p =0.0025 and ****p< 0.0001 (one-way ANOVA with Tukey’s post-hoc correction). N = 7, 8, 7, 7 with n = wells analyzed, three biological repeats. Scale bars: 20 µm. (B) Fluorescence lifetime images of Hilyte™ Fluor 488 labelled Aβ42 at the beginning of the experiment (t0) and after 2 h of incubation at room temperature (t2h), showing a decrease in the Aβ42 Hilyte™ Fluor 488 fluorescence lifetime when UCF-101 treated mitochondria were added. Data are presented as mean ± SEM. ***p =0.0009 and *p= 0.0142 (one-way ANOVA with Tukey’s post-hoc correction). N = 8, 8, 7, 8 with n = wells analyzed, three biological repeats. Scale bars: 20 µm.

### Inhibition of mitochondrial protein import enhances alpha-synuclein and Amyloid β 1-42 pathology

The effect of mitochondrial proteases on alpha-synuclein and Aβ42 aggregation could just lead to a general mitochondrial dysfunction, however a new line of thinking emerged recently, showing that mitochondrial proteases can influence aggregate dissolution and that aggregation-prone proteins are directed to mitochondrial import (42). There is one report showing mitochondrial import of alpha-synuclein (47), but it is discussed critically that aggregation-prone proteins like alpha-synuclein and Aβ42 are directly imported into mitochondria (48, 49). Thus, in order to prove whether we see alpha-synuclein residing within mitochondria, we immuno-gold labelled YFP-alpha-synuclein in SH-SY5Y cells, and found specific staining within mitochondria, which was mainly located at the inner mitochondrial membrane (Fig. 6A and Supplementary Fig. 2). In addition, we isolated mitochondria from wild-type adult rat brain and probed them for the presence of endogenous alpha-synuclein after protein K (PK) digestion. We saw that alpha-synuclein was still present after PK treatment, indicating that alpha-synuclein resides within the organelle since PK is not able to degrade proteins protected by organelle membranes. This is further supported, since incubation with 0.1% Triton X-100 during PK treatment, which is capable to solubilize mitochondrial membranes (50), enabled complete alpha-synuclein degradation (Fig. 6B and C).

**Fig. 6.**
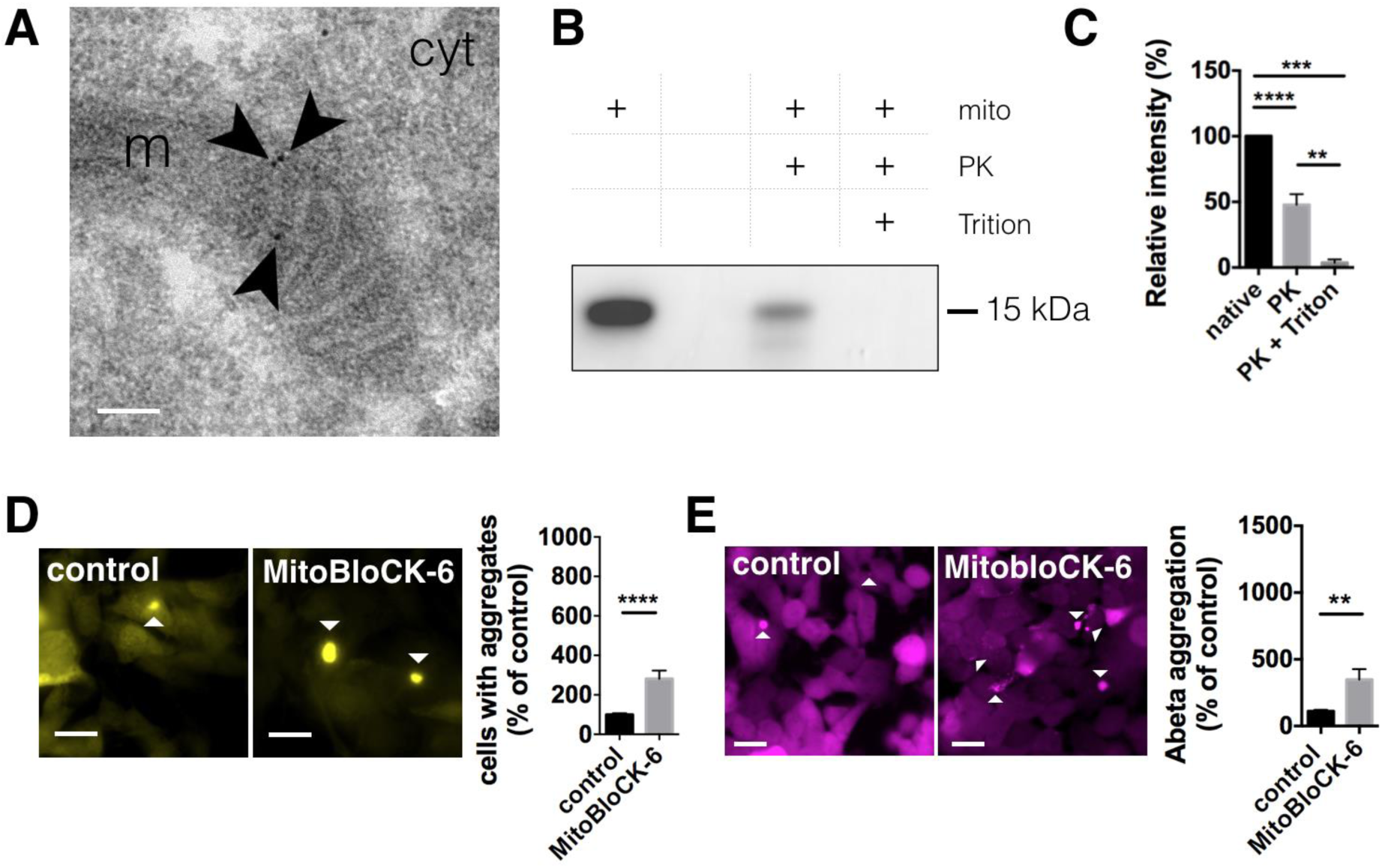
Inhibition of mitochondrial protein import increases alpha-synuclein and Aβ42 pathology. (A) Transmission electron microscopy (TEM) image of immunogold labelled YFP-alpha-synuclein in SH-SY5Y cells showing that alpha-synuclein is contained within mitochondria. Arrows indicate individual immunogold labels within mitochondria. m = mitochondria, cyt = cytoplasm. Scale bar: 100 nm. (B) Alpha-synuclein of isolated mitochondria from adult rat brain in the absence (native) of proteinase K (PK), in the presence of PK or in the presence of both PK and 0.1% TritonX-100. (C) Relative intensity of alpha-synuclein bands normalized to native mitochondria. Data are presented as mean ± SEM. ***p = 0.0007, ****p = < 0.0001, **p = 0.0017 (one-way ANOVA with Tukey’s post-hoc correction). N = 3 for all conditions with n = biological repeats. (D) YFP-alpha-synuclein SH-SY5Y were cells treated with DMSO (control), or 50 µM MitobloCK-6 before and during the incubation with alpha-synuclein fibrillar seeds (5h). Scale bars: 20 µm. Alpha-synuclein seeding was significantly increased upon treatment. Data are presented as mean ± SEM. ****p < 0.0001 (two-tailed Mann-Whitney U test). N = 15, 11 with n = regions analyzed, three biological repeats. (E) Aβ42-mCherry cells were treated with DMSO (control) or 5 µM MitobloCK-6 for 24 h. The aggregation of Aβ42 was increased upon treatment with MitobloCK-6. Data are presented as mean ± SEM. **p = 0.0088 (two-tailed unpaired t-test). N = 9, for all conditions, with n = wells analyzed, three biological repeats.

Since the above results indicate that alpha-synuclein is localized to mitochondria, we hypothesized that inhibition of mitochondrial protein import might have a similar effect on alpha-synuclein pathology as the inhibition of proteases. Thus, using MitobloCK-6, a small molecule inhibitor of protein translocation into mitochondria (51), we equally observed increased alpha-synuclein seeding in YFP-alpha-synuclein SH-SY5Y cells (Fig. 6D). Testing MitobloCK-6 on Aβ42-mCherry overexpressing HEK cells also showed increased aggregation (Fig. 6E), demonstrating that mitochondrial protein import influences the proteostasis of amyloidogenic proteins.

## Discussion

The effect of BAPTA-AM on alpha-synuclein seeding took us to study the role of mitochondrial proteostasis on the aggregation of amyloidogenic proteins. We demonstrate here, that inhibition of the mitochondrial proteases HtrA2 and Lon, as well as inhibition of mitochondrial protein import enhances alpha-synuclein pathology. However, downstream effects of mitochondrial dysfunction, induced without mitochondrial integrity disturbance, did not recapitulate the increased seeding. Inhibition of HtrA2 as well as inhibition of mitochondrial protein import also increased Aβ42 pathology, and the overexpression of HtrA2 was able to decrease Aβ42 aggregation notably. It was reported recently that mitochondria are able to influence the degradation and protein homeostasis of cytosolic proteins, which was shown in yeast cells upon heat shock (42), however mitochondria may also play an important role for the degradation of amyloidogenic proteins, since mitochondrial proteostasis seems to be clearly coupled to the pathology of alpha-synuclein and Aβ42. HtrA2 appears of particular interest, since it has previously been linked to PD by genetics (52–56) and shows a neuroprotective effect upon overexpression in mice (57, 58).

So far the effect of amyloid proteins on mitochondria has been interpreted only as a secondary pathological hallmark, with alpha-synuclein as well as Aβ exacerbating mitochondrial dysfunction (48, 49, 59–63). But amyloidogenic proteins might be directed to mitochondrial uptake deliberately, which then disrupts overall mitochondrial function if the uptake is overloaded. So vice versa, an initial failure in mitochondrial function, i.e. by complex I inhibition or upon disturbance of mitophagy, could eventually lead to increased levels of alpha-synuclein, having important implications for sporadic forms of the disease. Our recently published review provides more insights on the mitochondrial uptake of alpha-synuclein and Aβ, on the interaction with mitochondrial translocases as well as background on mitochondrial proteases (64).

There still remains the argument that inhibition of mitochondrial proteases just causes an unspecific mitochondrial dysfunction which then per se leads to increased alpha-synuclein aggregation. However, it seems that this effect is not mediated via the known downstream events of mitochondrial dysfunction. So it has been shown that ATP depletion is not able to increase alpha-synuclein aggregation, as seen in our study using short term complex I inhibition via MPP^+^ reducing ATP without causing major mitochondrial fragmentation and also previously when ATP levels were reduced independently from mitochondrial respiration using 2-Desoxyglucose, which inhibits glycolysis (65). Also increased calcium concentrations, when induced acutely via calcium influx through the plasma membrane using the ionophore ionomycin, did not influence alpha-synuclein seeding. This seems to stand in contrast to our previous study (25), where we have shown that calcium affects alpha-synuclein aggregation in-vitro. However, here mainly the nucleation rate was increased, thus calcium may still contribute to PD via alpha-synuclein seed formation. Oxidative stress has been discussed as a likely mechanism, since antioxidants are able to reduce dopaminergic neuron death and alpha-synuclein accumulation after complex I inhibition (65), however also a general protective impact on mitochondria might play a role.

Though, this does not mean that complex I inhibition, calcium dysregulation and oxidative stress do not play an important role during the course of the disease. Chronic complex I inhibition has clearly been shown to lead to dopaminergic neuron death and alpha-synuclein accumulation (12–16) and is a major factor implicating mitochondrial dysfunction in sporadic Parkinson’s disease. Chronic complex I inhibition would lead to reduced mitochondrial fitness and in a similar way high calcium loads do when specific neuronal subtypes are subjected to increased calcium concentrations over long time periods as shown recently for dopaminergic neurons of the substantia nigra (66). Taken together, our study shows that mitochondrial proteostasis may be an important factor in triggering pathology in neurodegeneration and attacking mitochondrial fitness, rather than downstream events of mitochondrial dysfunction might be crucial in the search for therapeutic strategies.

## Material and Methods

### Human cell culture

*Human neuroblastoma cells* (SH-SY5Y) were obtained from the European Collection of Cell Cultures (ECACC, Sigma-Aldrich, Dorset, UK) and grown in a 1:1 minimal essential medium (MEM) (Sigma-Aldrich) and nutrient mixture F-12 Ham (Sigma-Aldrich) supplemented with 15 % FBS, 1 % non-essential amino acids, 2 mM GlutaMAX and 1 % antibiotic-antimycotic (all Thermo Fisher Scientific, Epsom, UK). SH-SY5Y cells stably expressing YFP-alpha-synuclein were obtained by lentiviral transfection using 3^rd^ generation lentiviruses (Addgene constructs: 12251, 12253, 12259) (67). Human wild-type alpha-synuclein was inserted into EYFP plasmid (pEYFP-N1) using a 5 amino acid linker (sequence: GCACCGGTCGCCACC) between the C-terminus of alpha-synuclein and N-terminal EYFP. Alpha-synuclein-EYFP was then cloned into the pLJM1 backbone for lentiviral expression (Addgene: 19319) (68). For the preformed fibril (PFF) assay 50,000 cells were plated in MatTek dishes (P35G-1.5-14-C, MatTek Corporation, Ashland, US). For analysis of mitochondrial fragmentation cells were plated at 20,000 per well in NuncTM Lab-TekTM II Chambered Coverglass (8 well, 155409, Thermo Fisher Scientific).

Flp-InTM T-RExTM 293 cell line (Invitrogen), a derivative of HEK293 cells containing a stably integrated FRT site and a TetR repressor, were used to generate stable cell lines expressing either mCherry or Aβ42-mCherry (pcDNA3.3-mCherry, pcDNA3.3-Ab42-mCherry) under the Flp-InTM expression vector as described previously (44, 69). Cells were maintained in DMEM high glucose media (Sigma-Aldrich) supplemented with 10% fetal bovine serum (FBS), 2 mM glutaMAX, and 1 % antibiotic-antimycotic (all Thermo Fisher Scientific). Cells were grown at 37°C under a 5% CO2 atmosphere. Cells were plated at 35 000 cells per well in NUNC 24 well plates, and construct expression was induced for 3 days using media above with 1 µg/mL tetracycline (Sigma Aldrich) added. All cell lines were tested for mycoplasma contamination using the MycoAlert^TM^ PLUS mycoplasma detection kit (Lonza, Walkersville). For transient transfection of HtrA2 (70) electroporation with the NEON transfection system was used (settings: 1050 V, 30 ms, 2 pulses; Thermo Fisher Scientific). pcDNA3-HtrA2-FLAG was a gift from L. Miguel Martins (Addgene plasmid # 15938; http://n2t.net/addgene:15938; RRID:Addgene_15938).

Cells were imaged on a widefield microscope with IX83 frame (Olympus, Tokyo, Japan), HPLS343 plasma light source (Thorlabs, Newton, US), and Clara interline CCD camera (Andor, Belfast, UK), controlled by Micromanager (71). Respective filter cubes for YFP (excitation 500 nm, dichroic mirror 515 nm, emission 535 nm), RFP (excitation 560 nm, dichroic mirror 585 nm, emission 630 nm) and DAPI (excitation 350 nm, dichroic mirror 400 nm, emission 460 nm) were used. Images for YFP-alpha-synuclein aggregation and DAPI were taken with an Olympus Plan Apo U 60x/1.42 oil objective lens. Imaging was done randomly by automated acquisition of a grid of 7×7 images per area. Aggregates were identified by their fibrillar nature, cell nuclei were counted using FIJI (72). For Aβ42-mCherry aggregation images were taken with an Olympus LUCPlanFLN 20x/0.45 air objective lens. Aggregates were identified using the Thresholder plugin in ICY (73). The cell surface area was evaluated using the HK-Means plugin for ICY (74).

### Alpha-synuclein fibrils

Human wild-type (WT) alpha-synuclein was expressed in Escherichia coli One Shot® BL21 STAR™ (DE3) (Invitrogen, Thermo Fisher Scientific) cells using plasmid pT7-7 and purified using ion-exchange on a HiPrep Q FF 16/10 anion exchange column (GE Healthcare, Uppsala, Sweden) (75). Alpha-synuclein was then further purified on a HiPrep Phenyl FF 16/10 (High Sub) hydrophobic interaction column (GE Healthcare) (76). Purification was performed on an ÄKTA Pure (GE Healthcare). Monomeric protein was dialyzed against 20 mM phosphate buffer pH 7.2, lyophilized in a LyoQuest 85 freeze-dryer (Telstar, Spain), and stored at −80 °C.

Alpha-synuclein fibrils were produced by diluting alpha-synuclein monomer solution to a concentration of 150 µM in 20 mM phosphate buffer, pH 7.2. Samples were incubated at 37°C for 5 days in 0.5 mL Protein Lobind tubes (Eppendorf, Hamburg, Germany) under continuous rotation at maximum speed (UVP HB-1000 Hybridizer, Fisher Scientific). Fibrils were diluted 1:1 with 20 mM phosphate buffer, pH 7.2 to a final volume of 200 µL and sonicated (Digital Sonifier® SLPe, model 4C15, Branson, Danbury, USA) with six 10 sec pulses at 70 % amplitude and 10-sec pause after each sonication pulse. Sonicated fibrils were aliquoted, exposed to UV light for 30 min and frozen immediately after at −80C.

Alpha-synuclein fibrils were imaged by atomic force microscopy (AFM) (BioScope Catalyst microscope, Bruker AXS GmbH, Fitchburg, USA). Fibrils at an equivalent monomer concentration of 5 µM were deposited for 30 min on High-Performance cover glass (PN 474030-9020-000, Carl Zeiss Ltd.), cleaned for 30 min with 1 M KOH (Fluka, Bucharest, Romania) and coated for 30 min with 0.01 % poly-L-Lysine beforehand (P4707, Sigma). Samples were rinsed 5 times with deionized water and dried under nitrogen flow. AFM data were acquired using PeakForce Quantitative Nanomechanical Property mapping mode with ScanAsyst-Fluid+ probes (Bruker AXS GmbH). Images were flattened and exported using NanoScope Analysis software, version 1.8.

### Preformed fibril (PFF) assay

For the induction of alpha-synuclein seeding, YFP-alpha-synuclein overexpressing SH-SY5Y cells were incubated with sonicated preformed alpha-synuclein fibrils as described by Luk et al. (26). Briefly, cells plated in MatTek dishes were washed with Neurobasal medium and subsequently changed to 500 µL Neurobasal medium supplemented with 2 % B27 and 0.5 mM GlutaMAX (all Thermo Fisher Scientific). Cells were preincubated for 1 hour, either using 0.2 % DMSO for control or the respective treatment (see cell treatments below). 8 µL of PFFs were diluted with 32 µL HBSS (HBSS minus calcium and magnesium, no phenol red, 14175-053, Thermo Fisher Scientific) and mixed briefly 5 times. Fibrils were added to the bottom of the BioPORTER tube (BioPORTER® Protein Delivery agent, BP502424, Gelantis, San Diego, USA), mixed 5 times and incubated for 5 min at room temperature, then vortexed for 5 sec at 600 rpm (Stuart^TM^ Scientific SA8 vortex mixer, Sigma-Aldrich). 460 µL OptiMEM medium (Thermo Fisher Scientific) was added to the BioPORTER tube plus the respective treatments and mixed 5 times. The PFF mixture was added dropwise to the cells, settled and then incubated for 4 hours at 37°C and 5 % CO2. Final monomer equivalent concentration of preformed fibrils was 600 nM.

After 4 hours cells were washed twice with 1 ml Neurobasal medium and changed subsequently to 2 mL of retinoic acid medium made of 1:1 minimal essential medium (MEM) (Sigma-Aldrich) and nutrient mixture F-12 Ham (Sigma-Aldrich) supplemented with 5 % FBS, 1 % non-essential amino acids, 2 mM GlutaMAX and 1 % antibiotic-antimycotic (all Thermo Fisher Scientific) and 1 µM retinoic acid (Sigma-Aldrich) plus treatments if indicated and incubated for another 3 days to allow aggregate formation. Cells were fixed for 10 min using 4 % formaldehyde in PBS supplemented with 4 % sucrose, 5 mM MgCl2 and 10 mM EGTA, pH 7.4 (77), stained with Hoechst 33342 (Molecular Probes, Thermo Fisher Scientific) 1:2000 in PBS for 30 min.

### Cell treatments

Chemical used for the treatment of cells were prepared as followed, with final dilution made with the respective culture medium. Carbonyl cyanide 4-(trifluoromethoxy)phenylhydrazone (FCCP, Abcam, Cambridge, UK) 1 mM in DMSO, N-Methyl-4-phenylpyridinium Iodide (MPP+, Sigma-Aldrich) 10 mM in water, ionomycin (ab120370, Abcam) 10 mM and 1 mM in DMSO, 2-Deoxyglucose (Sigma-Aldrich) 0.5 M in water, menadione (Sigma-Aldrich) 1.5 mM in DMSO, BAPTA-AM (ab120503, Abcam) 2.5 mM in DMSO, BAPTA (ab144924, Abcam) 1 mM in water, CDDO-Me (Sigma-Aldrich) 1 mM in DMSO, UCF-101 (Sigma-Aldrich) 10 mM in DMSO and MitobloCK-6 (Focus Biomolecules) 5 mM in DMSO.

### Immunofluorescence

Cells were fixed as described above, blocking and permeabilisation were performed using 5 % donkey serum in 0.05 % Tween-20 in phosphate-buffered saline (PBS) for 1 h. Primary antibodies were incubated overnight at 4°C, followed by 5 washes with PBS. Secondary antibodies were incubated for 1 hour at room temperature, followed by 5 washes with PBS. As primary antibodies anti-Ubiquitin antibody, clone Apu2 (05-1307, 1:200, Millipore, Watford, United Kingdom), anti-Ubiquitin-binding protein p62, clone 2C11 (SQSTM1, 1:200, Abnova, Taipei, Taiwan) and anti-FLAG® M2 antibody (F1804, 1:200, Sigma-Aldrich) were used. As secondary antibodies anti-rabbit and anti-mouse Alexa Fluor®647, and anti-mouse Alexa Fluor®568 (A-21245, A-21236 and A-11031 from life technologies) were used. Samples were kept in PBS containing 5 mM sodium azide (Sigma-Aldrich).

### Structured illumination microscopy (SIM)

Structured illumination images were collected on a custom-built Structured Illumination Microscopy (SIM) setup which has been described in detail (78). A 60×/1.2NA water immersion lens (UPLSAPO 60XW, Olympus) focused the structured illumination pattern onto the sample. This lens also captured the samples’ fluorescence emission light before imaging onto a sCMOS camera (C11440, Hamamatsu). Laser excitation wavelengths used were 488 nm (iBEAM-SMART-488, Toptica), 561 nm (OBIS 561, Coherent), and 640 nm (MLD 640, Cobolt). Respective emission filters were BA 510-550 (Olympus), BrightLine FF01-600/37, and BrightLine FF01-676/29 (Semrock, New York, US). Imaging was done in fixed cells or live cells, as indicated. Images were acquired using custom SIM software (HCImage, Mamamatsu Corporation, Sewickley, US). Nine raw images were collected at each plane and each colour. FairSIM plugin in FIJI was used to reconstruct images (79).

### FLIM measurements of cytosolic calcium, H2O2, and ATP

Fluorescence lifetime microscopy (FLIM) was carried out on a custom-built Time-Correlated Single Photon Counting (TCSPC) system using a super-continuum laser (SC450, Fianium) with a pulse repetition rate of 40 MHz, a confocal scanning unit (FluoView 300, Olympus) coupled with an inverted microscope frame (IX70, Olympus), and a time-correlated single-photon counting system (Becker & Hickl GmbH) as described in detail before (80). The excitation wavelength was selected by using an acousto-optic tunable filter (AOTFnC-400.650, Quanta Tech) and respective excitation filters (to improve the wavelength selection), and emission fluorescence was imaged through respective emission filters. The data acquisition time was 200s for each FLIM image (10 cycles, 20s per cycle). The photon detection rate was kept below 2% of the laser repetition rate in order to avoid photon pile-up.

For cytosolic calcium measurements SH-SY5Y cells were incubated with Oregon Green^TM^ 488 BAPTA-1, AM (Thermo Fisher Scientific) for 45 min at 1 µM concentration. Excitation was set to 475 nm, excitation filter BrightLine FF01-474/27 (Semrock), and emission filter BrightLine FF01-525/39 (Semrock) were used. For measurement of H2O2 and ATP SH-SY5Y cells were transiently transfected with the respective sensor using electroporation with the NEON transfection system (settings: 1100 V, 50 ms, 1 pulse; Thermo Fisher Scientific). HyPer, a genetically encoded sensor consisting of circularly permuted yellow fluorescent protein (cpYFP) inserted into the regulatory domain of the prokaryotic H2O2-sensing protein, OxyR (81) was used to measure cytosolic hydrogen peroxide. Excitation was set to 470 nm, same excitation and emission filters as for Oregon Green^TM^ 488 BAPTA-1 were used. Ateam1.03, a FRET-based indicator for ATP composed of the ε subunit of the bacterial FoF1-ATP synthase sandwiched by CFP and YFP (82, 83) was used to measure cytosolic ATP levels. Excitation was set to 435 nm, excitation filter BrightLine FF01-434/17 (Semrock), and emission filter BrightLine FF01-470/28 (Semrock) were used. ATeam1.03-nD/nA/pcDNA3 was a gift from Takeharu Nagai (Addgene plasmid # 51958; http://n2t.net/addgene:51958; RRID:Addgene_51958). For ATP measurements cells were subjected to media containing 10 mM 2-Deoxyglucose to inhibit glycolysis. The fluorescence lifetime was analyzed by the FLIMfit software tool developed at Imperial College London (84, 85).

### ThT Assay

The aggregation of alpha-synuclein in vitro was measured by Thioflavin T (ThT) assay. Briefly, 50 µL of 100 µM alpha-synuclein with 10 µM fresh ThT added, was incubated for 7 days with 1% DMSO as a control, 10 µM FCCP, 10 µM BAPTA-AM, or 10 µM BAPTA. Assays were performed in NUNC™ black 384-well plates with optical flat bottom (142761, Thermo Fisher Scientific) which were sealed with an Ampliseal transparent microplate sealer (Greiner Bio-One GmbH). Plates were incubated with orbital shaking at 300 rpm for 5 minutes before each read every hour at 37 °C for 170 cycles. The readings of ThT fluorescence intensity were taken using excitation at 440 nm and emission at 480 nm, collected from the bottom up with 20 flashes per well and a gain setting of 1300 (FLUOstar Omega, BMG Labtec GmbH, Ortenberg, Germany). Experiments were repeated three times with four replicates for each condition.

### Mitochondrial fragmentation

To label mitochondria SH-SY5Y cells were incubated overnight with 1:1000 CellLight^TM^ Mitochondria-RFP (Thermo Fisher Scientific) and imaged by SIM or a widefield microscope for quantification. Images were taken randomly by automated imaging of a grid, images were analyzed from 3 biological repeats. The mitochondrial length was evaluated using the NIEL Mito algorithm (86, 87).

### Animals

Adult female Sprague Dawley rats were supplied by Charles River UK Ltd., Scientific, Breeding and Supplying Establishment, registered under Animals (Scientific Procedures) Act 1986, and AAALAC International accredited. All animal work conformed to guidelines of animal husbandry as provided by the UK Home Office. Animals were sacrificed under schedule 1; procedures that do not require specific Home Office approval. Animal work was approved by the NACWO and University of Cambridge Ethics Board.

### Mitochondrial isolation and Western blot analysis

Mitochondria were isolated from adult rat brain by differential centrifugation using mitochondria isolation kit for tissue (ab110168, abcam). Western blot for alpha-synuclein was performed using 4–12% Bis-Tris gels (Life Technologies), the protein was transferred onto 0.45 µm Millipore PVDF membrane (Fisher Scientific, Loughborough, UK) and subsequently fixed using 4% formaldehyde + 0.1% glutaraldehyde in PBS (both Sigma-Aldrich) (88). As primary antibody α-Synuclein (D37A6) XP® Rabbit mAb was used (1:1000 dilution, #4179, CST, Leiden, Netherlands). An enhanced chemiluminescence (ECL)-horse radish peroxidase (HRP) conjugated secondary antibody (NA934V, 1:1000 dilution, GE Healthcare, Uppsala, Sweden) and SuperSignal West Femto Chemiluminescent Substrate (Thermo Fisher Scientific) were used to probe the membrane, which was exposed using a G:BOX (Syngene, Cambridge, UK). Western blots were analyzed in FIJI (72).

### TEM

SH-SY5Y cells and SH-SY5Y cells overexpressing YFP-alpha-synuclein were cultured in 6 well plates (Greiner Bio-One GmbH) at 350 000 per well. After reaching confluency cells were washed with 0.9% NaCl (Sigma-Aldrich) twice and incubated with 8% formaldehyde in 0.05 M sodium cacodylate buffer (Paraformaldehyd from Merck, Darmstadt, Germany) pH 7.4 for 2h at 4°C. Cells were scraped from 6 wells and centrifuged for 10 min at 3500 g. Cells were washed 5 times in 0.05 M sodium cacodylate buffer, 3 times in deionized water, and incubated with 2 % uranyl acetate in 0.05 maleate buffer pH 5.2 (both BDH Chemicals Ltd., Dorset, UK) overnight at 4°C. Cells were washed again and dehydrated at increasing ethanol concentrations (1x 50% EtOH, 3x 70% EtOH, 3x 95 % EtOH, 3x 100% EtOH, 3x 100 % dry EtOH; 5 min in each, Sigma-Aldrich). Cells were resuspended in LRW resin (LR White Resin, London Resin (Hard), Agar Scientific, Stansted, UK) mixed 50/50 with dry 100% EtOH and incubated overnight at room temperature. The following day, cells were spun down, and resuspended in pure LRW for 2 days, where LRW was exchanged twice. Cells were centrifuged at 13000 g to form a firm pellet, which was transferred to size 2 gelatine embedding capsules (TAAB, Aldermaston, UK) containing LRW resin. Gelatine capsules were covered with a glass coverslip to exclude any air and the resin was cured at 60°C for 2 days. Gelatine capsules were removed and ultrathin sections were cut using a Leica Ultracut E Ultramicrotome (Leica, Wetzlar, Germany) and placed on 400 mesh nickel/formvar film grids (EM Resolutions). Sections were stained with Anti-GFP antibody (ab6556, Abcam) in blocking solution (2 % BSA (BBITM solutions, Crumlin, UK) in 10 mM TRIS (Sigma-Aldrich) buffer pH 7.4 containing 0.001% Triton-X100 (Calbiochem, San Diego, US) and 0.001% Tween20 (Sigma-Aldrich) at 1:100 overnight. After washing, sections were incubated with goat anti-rabbit 10 nm gold secondary antibody (BBITM solutions) in blocking solution at 1:200 for 1 hour. Sections were washed with washing buffer (same as above omitting BSA), deionized water and left for drying overnight. Post-staining included 2% uranyl acetate in 50 % methanol for 30 sec, followed by washing with 50 % methanol and 30-sec staining in Reynold’s lead citrate (lead nitrate from BDH Biochemicals Ltd., Trisodiumcitrate from Sigma-Aldrich). Grids were rinsed thoroughly with deionized water and dried before imaging. Grids were imaged on an FEI Tecnai G2 electron microscope (Thermo Fisher Scientific) run at 200 keV using a 20 µm objective aperture, images were taken using an AMT V600 camera (AMT, Woburn, US).

### In-vitro measurements of Aβ42 aggregation

Synthetic Aβ42 and Aβ42 Hilyte™ Fluor 488 (both from Anaspec, Seraing, Belgium) were prepared as previously described (89). Briefly, lyophilized Aβ42 (1 mg) was dissolved in ice-cold trifluroacetic acid (200 mL), sonicated at 0 °C for 60 s and then lyophilized overnight. Ice cold 1,1,1,3,3,3-hexafluro-2-propanol (1 mL) was added, sonicated at 0 °C for 60 s and aliquoted as 20 μL units. The samples were lyophilized overnight and were stored at −80 °C until use. Lyophilized Aβ42 Hilyte™ Fluor 488 peptide (0.1 mg) was dissolved in 1% NH4OH (200 μL) and sonicated for 60 s at 0 °C. The sample was aliquoted into 5 μL units, snap-frozen in liquid nitrogen and stored at −80 °C. Immediately before the experiment unlabeled Aβ42 was prepared by adding first dimethyl sulfoxide (DMSO) (5% of total solvent volume), then sodium phosphate buffer (NaP buffer 50mM, pH 7.4) to reach a concentration of 20 µM. The solution was sonicated at 0 °C for 3 min and centrifuged at 13,400 rpm at 0 °C for 30 min. Then the sample was further diluted to 5 µM concentration with NaP buffer. Also the labelled Aβ42 Hilyte™ Fluor 488 was brought to 5 µM concentration in NaP buffer and both were mixed in 1:1 ratio. Samples were prepared on ice adding Aβ42, 1 mg/mL of purified mitochondria (preparation see above) and 20 µM UCF-101. Mitochondria isolation buffer and DMSO were added in control samples. 12 µL volume were pipetted in silicon gaskets (Thermo Fisher Scientific, P24742) on a coverslip and measured at room temperature. Fluorescence lifetime measurements (FLIM) were carried out on a custom-built Time-Correlated Single Photon Counting (TCSPC) system as described above (see FLIM measurements of cytosolic calcium, H2O2, and ATP).

### Statistics

Statistical analysis was performed using GraphPad Prism 6.07 (GraphPad Software, Inc., La Jolla, CA, USA). Values are given as mean ± SEM unless otherwise stated. Normal distribution was tested using Shapiro-Wilk test. Two-tailed unpaired t-test was used upon normal distribution, two-tailed Mann-Whitney U test was used when no normal distribution was given. For multiple comparisons either one-way ANOVA with Dunnett’s post hoc correction upon normal distribution or Kruskal-Wallis test with Dunn’s multiple comparison when no normal distribution was given were performed. Significance was considered at p < 0.05.

## Data availability

All relevant data are available from the corresponding authors.

## Acknowledgements

We would like to thank Karin H. Muller and Jeremy N. Skepper for their help and input for the transmission electron microscopy study. We would like to thank Samantha Beck for establishing the YFP-alpha-synuclein SH-SY5Y cell line. J.L. was supported by a research fellowship from the Deutsche Forschungsgemeinschaft (DFG; award LA 3609/2-1). C.F.K. acknowledges funding from the UK Engineering and Physical Sciences Research Council (EPSRC). G.S.K.S. and C.F.K. acknowledge funding from the Wellcome Trust, the UK Medical Research Council (MRC), Alzheimer Research UK (ARUK), and Infinitus China Ltd. J.L. and A.D.S. acknowledge Alzheimer Research UK (ARUK) travel grants.

## Author contributions

J.L., A.F.V. and A.M. contributed to alpha-synuclein PFF assay and cell work. J.L performed FLIM experiments and mitochondrial morphology analysis. A.D.S. performed alpha-synuclein in vitro work. J.L. and A.F.V. and C.H. contributed to Western blot analyses. J.L. performed TEM. J.L., S.W.V. and M.L. contributed to Aβ42 studies. J.M. and M.F. contributed to automated and SIM imaging. J.L. designed the experiments. J.L., C.F.K. and G.S.K.S. conducted the overall manuscript.

## Figure Legends

**Supplementary Fig. 1.**
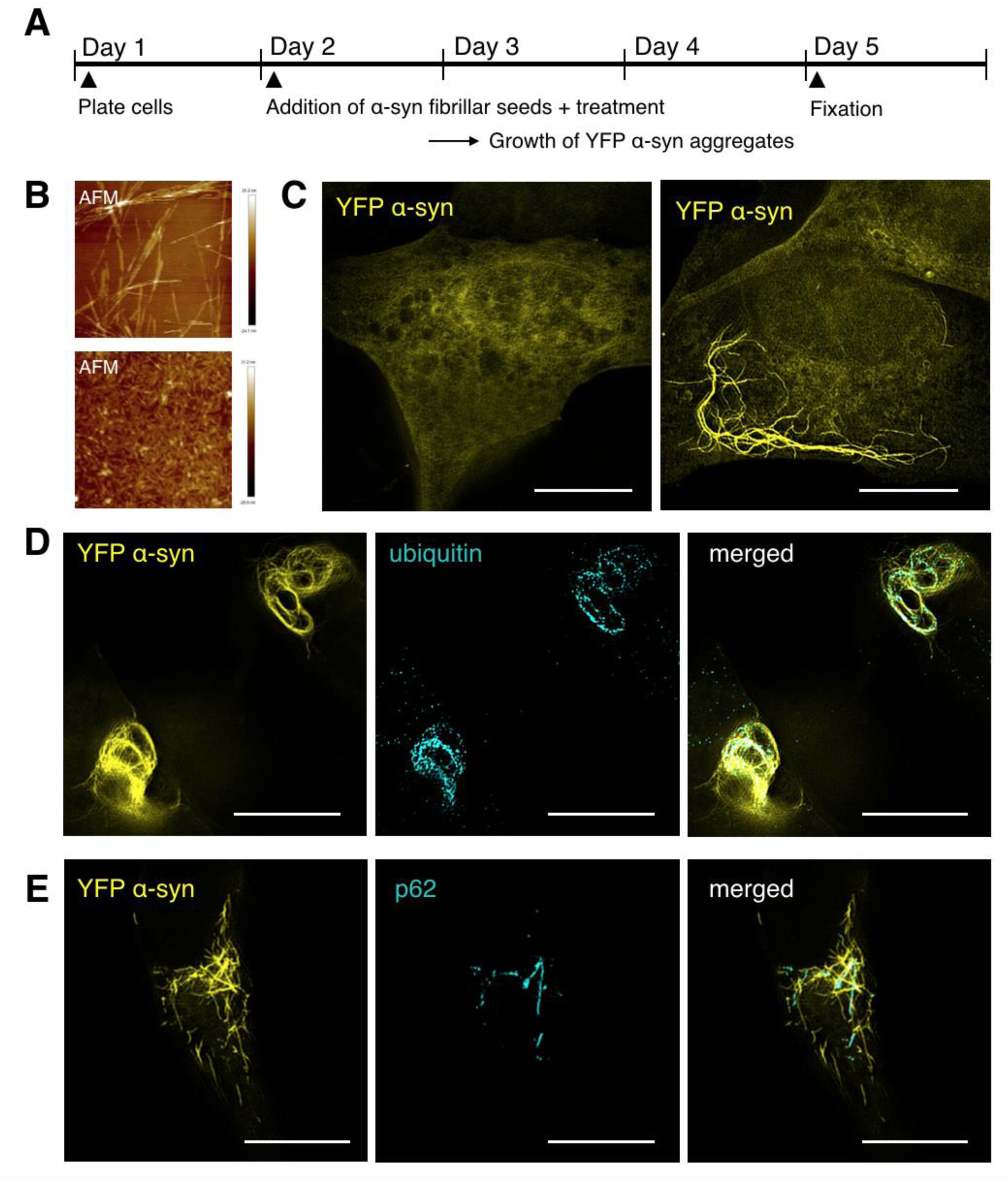
Incubation of SH-SY5Y cells overexpressing YFP-alpha-synuclein with alpha-synuclein seeds leads to YFP-alpha-synuclein fibril formation. (A) Schematic overview of the cellular alpha-synuclein aggregation assay. (B) Fibrillar seeds generated from recombinant human wild-type alpha-synuclein shown by atomic force microscopy before (upper panel) and after sonication (lower panel). Scale bars: 1 µm. (C) Structured illumination microscopy (SIM) images of SH-SY5Y cells overexpressing YFP-tagged alpha-synuclein in the absence (left) and upon incubation with alpha-synuclein fibrillar seeds (right). Scale bars: 10 µm. (D and E) Co-staining of YFP-alpha-synuclein fibrillar aggregates with ubiquitin and ubiquitin-binding protein p62. Scale bars: 10 µm.

**Supplementary Fig. 2.**
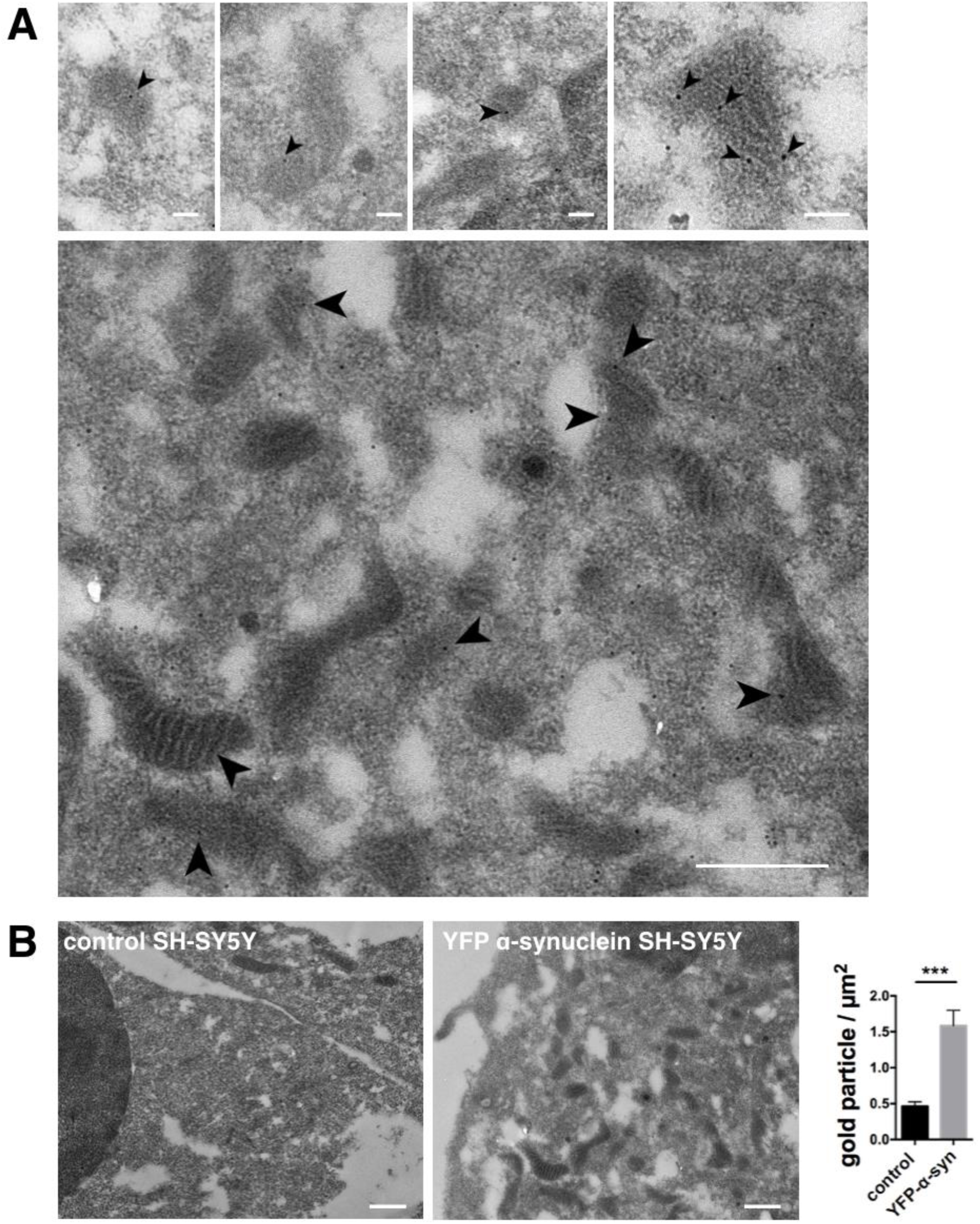
TEM of immunogold labelled YFP-alpha-synuclein, supplementary to Fig. 6A. (A) Representative images of transmission electron microscopy (TEM) from SH-SY5Y cells overexpressing YFP-alpha-synuclein showing that alpha-synuclein is contained within mitochondria. Arrows indicate individual immunogold labelling within mitochondria. Scale bars: 100 nm small images, 500 nm large overview image. (B) TEM images and quantification of anti-GFP staining in control SH-SY5Y cells and SH-SY5Y cells overexpressing YFP-alpha-synuclein. Quantification of gold particles / µm^2^ shows that the staining is enriched in YFP-alpha-synuclein overexpressing cells and not due to unspecific background. Data are presented as mean ± SEM. ***p = 0.0002 (two-tailed unpaired t-test). N = 10, 13 with n = images analyzed. Scale bars: 500 nm.

